# Full-featured, real-time database searching platform enables fast and accurate multiplexed quantitative proteomics

**DOI:** 10.1101/668533

**Authors:** Devin K. Schweppe, Jimmy K. Eng, Derek Bailey, Ramin Rad, Qing Yu, Jose Navarrete-Perea, Edward L. Huttlin, Brian K. Erickson, Joao A. Paulo, Steven P. Gygi

**Affiliations:** Department of Cell Biology, Harvard Medical School, Cambridge, MA; University of Washington Proteomics Resource, Seattle, WA; Thermo Scientific LSMS, San Jose, CA

## Abstract

Multiplexed quantitative analyses of complex proteomes enable deep biological insight. While a multitude of workflows have been developed for multiplexed analyses, the most quantitatively accurate method (SPS-MS3) suffers from long acquisition duty cycles. We built a new, real-time database search (RTS) platform, Orbiter, to combat the SPS-MS3 method’s longer duty cycles. RTS with Orbiter enables the elimination of SPS-MS3 scans if no peptide matches to a given spectrum. With Orbiter’s online proteomic analytical pipeline, which includes RTS and false discovery rate analysis, it was possible to process a single spectrum database search in less than 10 milliseconds. The result is a fast, functional means to identify peptide spectral matches using Comet, filter these matches, and more efficiently quantify proteins of interest. Importantly, the use of Comet for peptide spectral matching allowed for a fully featured search, including analysis of post-translational modifications, with well-known and extensively validated scoring. These data could then be used to trigger subsequent scans in an adaptive and flexible manner. In this work we tested the utility of this adaptive data acquisition platform to improve the efficiency and accuracy of multiplexed quantitative experiments. We found that RTS enabled a 2-fold increase in mass spectrometric data acquisition efficiency. Orbiter’s RTS was able to quantify more than 8000 proteins across 10 proteomes in half the time of an SPS-MS3 analysis (18 hours for RTS, 36 hours for SPS-MS3).

## Introduction

Multiplexed quantitative methods continually attempt to balance acquisition speed and precursor isolation purity. The balance derives from the need to achieve high proteome coverage (speed, depth) to interrogate new biologies and the need for quantitative accuracy to eliminate spurious quantitative values (purity, accuracy) to improve quantitative dynamic range. Initial experiments with multiplexed isobaric reagents implemented HRMS2-based methods for multiplexed quantitation and relied on a single precursor isolation to attempt to eliminate co-isolating ions^1^. While the methods were relatively fast, the resulting quantitation suffered from the well-documented phenomenon of reporter ion interference due to co-isolation of precursors^2^.

In an effort to eliminate the aforementioned quantitative interference, methods that employed a tertiary scan to analyze secondary fragmentation products, multinotch MS3 or SPS-MS3, were developed^2,3^. The SPS-MS3 method vastly improved quantitative accuracy, but required the addition of a third quantification scan to every instrument scan cycle which subsequently slowed instrument acquisition speeds^3,4^. While other methods have been developed to reduce precursor co-isolation interference and/or increase the duty cycle speed, these methods generally still rely on either the HRMS2 method or SPS-MS3 method^5,6^. Recently a proof-of-principle showed real-time spectral matching as a novel means to achieve the speed of HRMS2 analyses with the quantitative accuracy of SPS-MS3^4^. Real-time search (RTS) had the potential to vastly improve acquisition efficiency for multiplexed quantitative analysis by enabling quantitation if and only if a peptide spectral match (PSM) was found^4^. This intelligent acquisition strategy was based around a binomial search score^7,8^. By applying this strategy specifically to multiplexed analyses and through selective elimination of SPS-MS3 this study demonstrated a strong improvement in scan acquisition speed and improved accuracy for multiplexed quantitation.

In the present work we extended the robustness and flexibility of the RTS strategy for multiplexed quantitative proteomics. We implemented a full analytical pipeline – monoisotopic peak refinement, database searching, and FDR filtering^9^ – on a millisecond time scale, termed Orbiter. The speed of Orbiter analysis enabled real-time decision making to dictate the acquisition of SPS-MS3 scans only when a PSM was observed. We chose the open source Comet search engine for database searching and scoring^10,11^. For this work, Comet was revised to enable fast real-time spectral analysis while maintaining support for highly flexible searching (e.g. post translation modifications, multiple isotopic envelopes, and flexible fragmentation schemes)^10,12^. We evaluated the improved performance of Orbiter RTS against standard SPS-MS3. Orbiter achieved 2-fold faster acquisition speeds and improved quantitative accuracy compared to canonical SPS-MS3 methods.

## Experimental Methods

### Tissue culture and sample preparation

Yeast cells (*Saccharomyces cerevisae*, BY4742) were grown in 500mL YPD cultures to an OD600 of 0.8 then washed twice with ice-cold PBS, pelleted, and stored at - 80° until use. Cells were resuspended in lysis buffer (8M urea, 50mM EPPS pH 8.5, 150mM NaCl, Roche protease inhibitor tablet) and lysed by bead beating. After lysis and bead removal, the lysate was centrifuged to remove cellular debris and the supernatant was collected for use. Cell lines were grown to confluence in DMEM containing 10% fetal bovine serum and 1% streptomycin/puromycin. Cells were harvested by manual scraping and washed twice with PBS. Cells were syringe lysed in lysis buffer (8M urea, 50mM EPPS pH 8.5, 150mM NaCl, and Roche protease inhibitor tablet) and the resulting lysates were cleared via centrifugation.

Desired protein amounts were aliquoted and chloroform methanol precipitated, followed by digestion with LysC (overnight at room temperature, vortex speed 2; Wako) and trypsin (6 hours, 37° Promega) digestion. Peptides were labeled with TMT reagents as previously described^6,13^. Labeled peptides were mixed, and dried to remove organic solvent prior to clean-up via Sep-Pak (50mg C18 SepPak; Waters). As needed, labeled peptide mixtures were separated via high-pH reversed phase chromatography and pooled into 12 fractions^13^. Samples were dried and stored at −80° prior to analysis.

### LC-MS/MS Analysis

Samples were resuspended in 5% acetonitrile/2% formic acid prior to being loaded onto an in-house pulled C18 (Thermo Accucore, 2.6A, 150um) 35cm column. Peptides were eluted over a 90, 120, or 180 minute gradient from 96% buffer A (5% acetonitrile, 0.125% formic acid) to 30% buffer B (95% acetonitrile, 0.125% formic acid). Sample eluate was electrosprayed (2600V) into a Thermo Scientific Orbitrap Fusion Lumos mass spectrometer for analysis. The scan procedure for MS1 scans (Orbitrap scan at 120,000 resolution, 50ms max injection time, and AGC set to 1e5) and MS2 scans (Rapid ion scan, 50ms max injection time, AGC set to 2e4, CID collision energy of 35% with 10ms activation time, and 0.5 m/z isolation width) was constant for all analyses.

### Database search and analysis

Raw files were converted to mzXML format using an in-house adapted version of RawFileReader^6^ and searched using SEQUEST or Comet^11,14^. Briefly, spectra were searched against a target-decoy database for the yeast, human, or concatenated human-yeast proteomes, including isoforms^6^. Searches were performed with a 20 ppm peptide mass tolerance, 0.9 Da fragment ion tolerance, trypsin enzymatic cleavage with up to 2 missed cleavages, and three variable modifications allowed per peptide. Unless otherwise noted, all searches were performed with variable methionine oxidation (+15.9949146221), static cysteine carboxyamido-methylation (+57.02146) and static tandem mass tag modifications on lysine and the peptide N-termini (+229.16293). Peptide spectral matches were filtered to a peptide and protein false discovery rate (FDR) of less than 1%^15^. Statistical analyses and plotting was done using the R project for statistical computing^16^.

### Adaptive instrument control

The adaptive instrument control platform (Orbiter) was built in the .NET Framework (v4.6.5). Peptide spectral matches were determined using a version of the Comet search algorithm specifically designed for improved spectral acquisition speed enabling searching full target-decoy databases. These improvements have been made available in the latest release of Comet^10,11^. Single spectrum searching via this modified revision of Comet retains the full complement of search features available to Comet (e.g. static/variable modifications, indexed databases) enabling highly customizable searches. The real-time search (RTS) Comet functionality has been released and is available here: http://comet-ms.sourceforge.net/. Real-time access to spectral data was enabled by the Thermo Scientific Fusion API (https://github.com/thermofisherlsms/iapi). The core search functionalities demonstrated here have been incorporated into the latest version of the Thermo Scientific instrument control software (Tune 3.3).

### Real-time monoisotopic peak correction

Monoisotopic peaks were corrected in real-time to attain highly accurate monoisotopic m/z values for each precursor via modeling averagine across a given precursor. Briefly, potential monoisotopic envelope peaks are extracted using an averagine mass offset^17^. Peaks are then compared using Pearson correlation against a theoretical distribution calculated from the estimated number of carbons and the natural abundance of ^13^C. The mass with the best correlation is used as the monoisotopic m/z^17^.

### Real-time false discovery rate estimation

To improve scoring for a potentially diverse cohort of samples, real-time false discovery rate (rtFDR) filtering was implemented using a modified linear discriminant analysis^15^ adapted from the Accord. Net Statistical libraries^18^. Seven parameters were used for rtFDR estimation: XCorr, deltaCorr, missed cleavages, charge state, absolute ppm error, peptide length, and the fraction of ions matched^15^. After requiring a minimum set of observed reverse hits, discriminant scores were used to sort and filter PSMs to a user-defined rtFDR^19^. Continuous real-time peptide spectral matching also enabled on-the-fly instrument ppm error estimation and correction.

## Results and Discussion

During mass spectrometric analyses, stochastic precursor selection and fragmentation results in large numbers of non-peptide matching spectra^4^. In standard SPS-MS3 methods, these non-matching spectra generate wasteful MS3 scans. We reasoned that elimination of these extraneous SPS-MS3 scans would enhance SPS-MS3 acquisition speed while maintaining quantitative accuracy^3,20^. Therefore, performing RTS prior to SPS-MS3 acquisition and only triggering SPS-MS3 scans when a PSM was observed could significantly reduce the number of spurious SPS-MS3 scans generated^4^. This process would free instrument time to acquire more peptide matching MS2 scans to increase proteome coverage which would in turn trigger useful SPS-MS3 scans. We built the Orbiter platform to perform this RTS decision making. Orbiter encompassed a full featured analytical proteomics pipeline operating fast enough to seamlessly integrate concurrently with instrument scan acquisition and inform future scan decisions (i.e. when to trigger SPS-MS3).

The Orbiter platform was built in C# (.NET 4.6.5+) to perform spectral pre-processing, RTS using the Comet search engine, SPS ion selection, and real-time false discovery rate filtering within milliseconds of the MS2 scan acquisition (Figure 1). Three central components enabled fast, adaptive database searching. First, the processing of precursor MS1 scans prior to database searching facilitated monoisotopic precursor mass assignment and correction on the fly. Second, a newly developed revision of the Comet database searching algorithm enabled rapid single spectrum searching. Third, real-time false discovery rate (FDR) filtering was built using linear discriminant analysis to filter PSMs using user-defined FDR settings.

**Figure 1.**
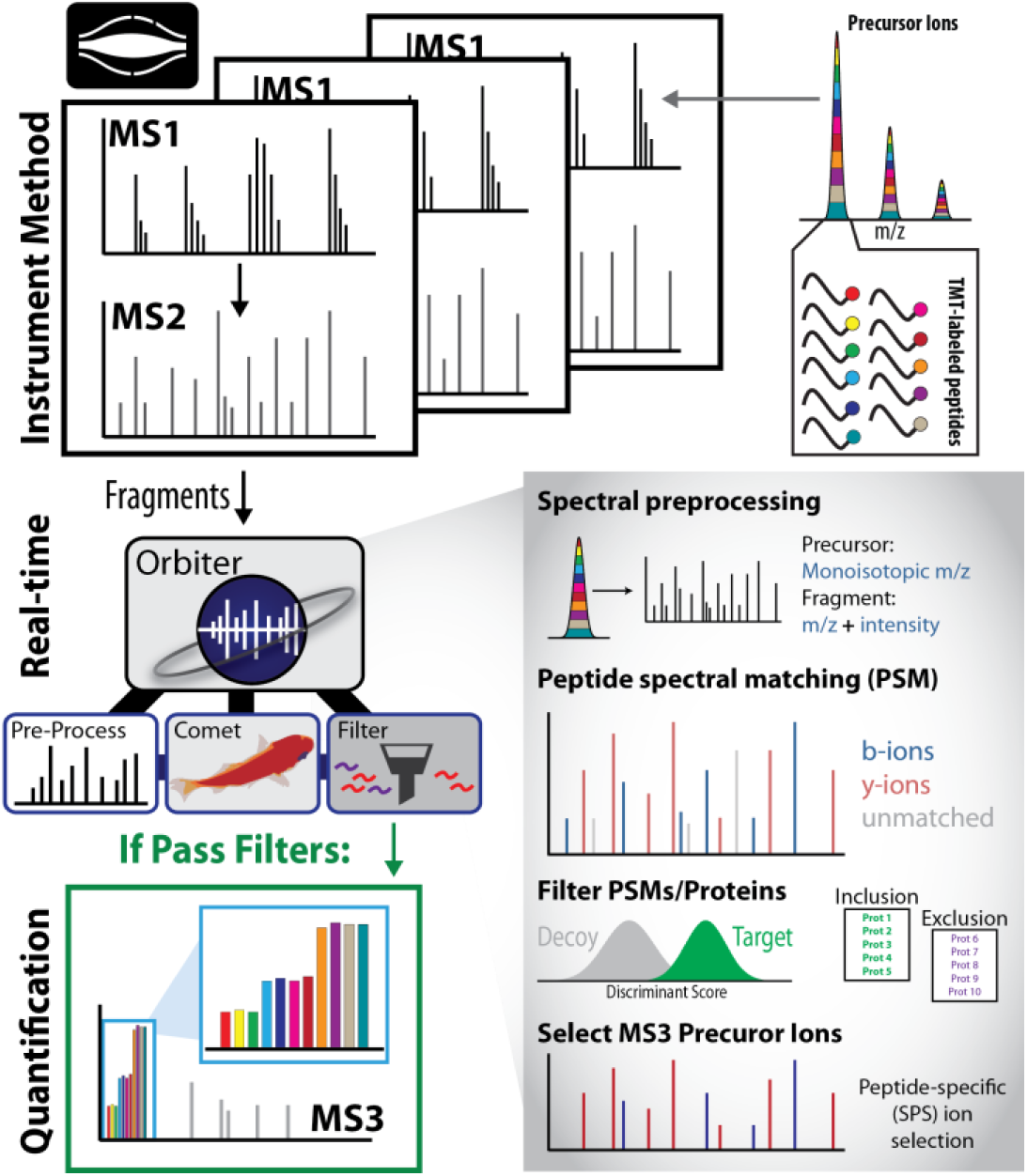
Architecture of the Orbiter platform. DDA MS2 scans are searched by Orbiter’s real-time database search. Peptide spectral matches are identified using Comet and filtered using a real-time FDR filter. For multiplexed analyses, SPS ions based on b- and y-ions from the identified peptide sequence. If and only if a peptide spectral match passes all assigned filters, will Orbiter trigger a new SPS-MS3 scan.

To run an RTS analytical method, a user queues a canonical DDA MS2 acquisition method (MS1 precursors trigger MS2 fragmentation scans for ion trap or Orbitrap analysis) based on user specified filters (e.g. dynamic exclusion, intensity thresholds). Running on the instrument computer, Orbiter listens for the beginning of each new method. When a new method begins, Orbiter then listens for the scan description of the first three new MS1 scans. If the consensus scan description matches to a previously established Orbiter method, Orbiter will begin searching with the pre-established database and parameters. Otherwise, users can override Orbiter methods by inputting a specific indexed database and parameters. Instrument listening and control was accomplished using Thermo Scientific’s Tribrid instrument application programming interface (iAPI).

Refinement of monoisotopic peak detection has previously been shown to improve peptide identification rates^17,21,22^. To ensure fidelity to the actual monoisotopic peak mass (often corrected offline) and to offline search workflows, we developed a fast monoisotopic peak algorithm to correct instrument assignments based on an averaged precursor monoisotopic mass^17^. Here, Orbiter collects the centroids for every MS1 scan in real-time and generates a library of all potential precursors that may be targeted for fragmentation by the instrument. Peaks that could potentially be part of the isotopic envelope were extracted using the averagine mass offset^17^. Peaks were then compared using Pearson correlation against a theoretical distribution calculated from the estimated number of carbons and the natural abundance of ^13^C. The resulting peak with the highest correlation was assigned as the monoisotopic peak. Finally, the monoisotopic peak detection averages the monoisotopic mass for the triggering precursor across the preceding *n* MS1 scans. This procedure enables accurate monoisotopic peak detection for use in subsequent searching of potential peptide fragments.

Comet was chosen as the database searching algorithm as it was built under an open source framework with active revisions and maintenance, allowing for rapid prototyping and adaptation to the challenges of real-time search^10,11^. Additionally, Comet has an extensive suite of multi-application search features^10^, including well documented and validated search and scoring functions – for example XCorr and deltaCorr^10,11^. The primary challenge for integrating Comet into an RTS engine was to streamline the scoring functions and optimize memory allocation to ensure Comet searching could be run efficiently^10^. We accomplished this by deploying new finalizers to rapidly expunge unmanaged memory resources. This enabled RTS to run stably over extended periods of time (weeks to months).

In addition, the RTS revision of Comet was adapted to remove post-search E-value calculations^11^. The E-value calculation made up greater than 80% (e.g. 125ms of a total of 156ms) of the Comet processing time. Thus, by eliminating these calculations, search speeds increased 5-fold, but required a new FDR metric (see below). With these changes, Orbiter was able to rapidly search each new MS2 spectra against an entire organismal database in milliseconds with high fidelity to the Comet run offline (Figure 2a). When comparing to offline searching using Thermo’s high throughput SEQUEST, we observed highly correlated scores similar to previous comparisons of the two search engines (Figure 2A)^10,14^.

**Figure 2.**
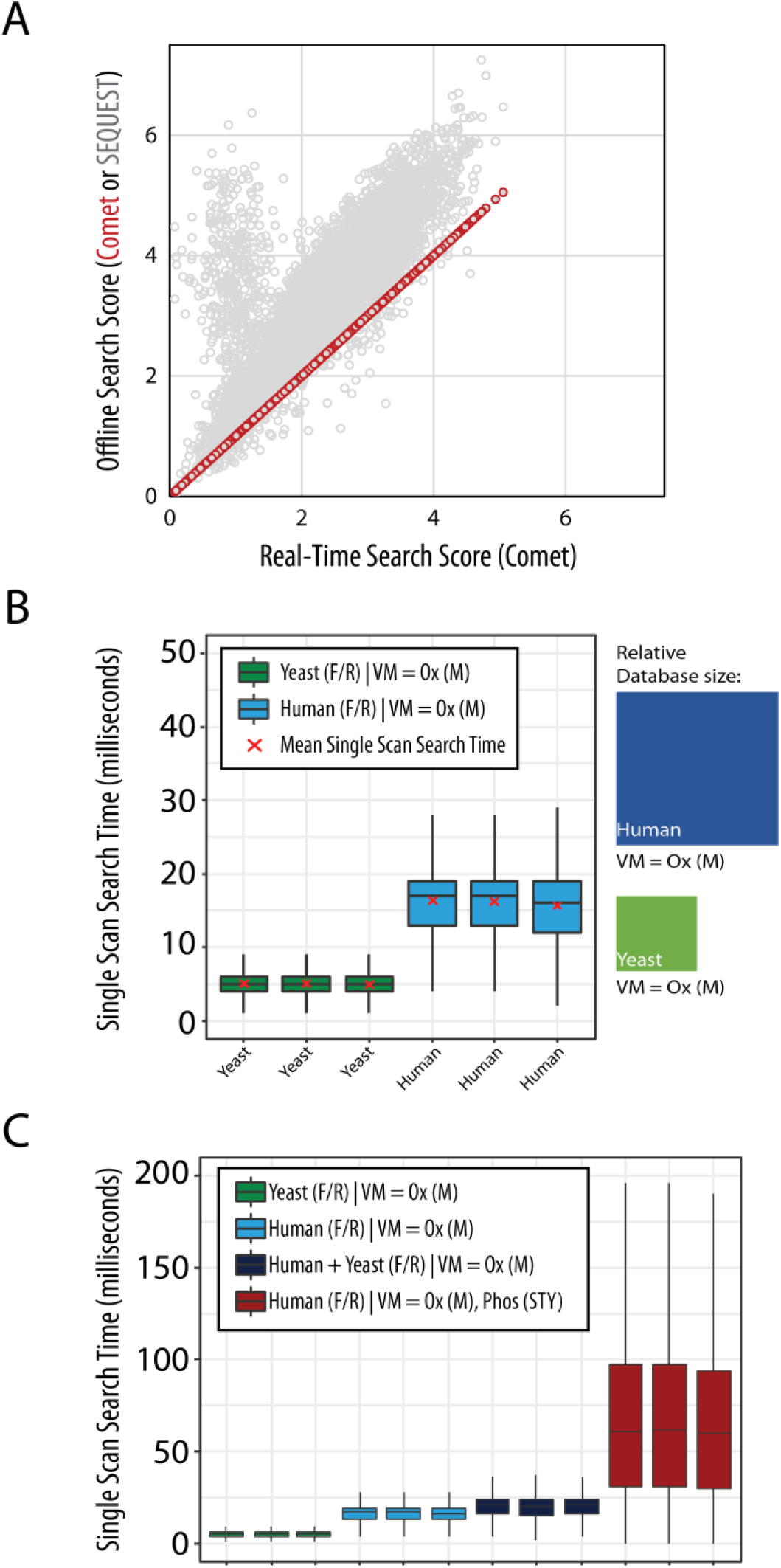
Real-time searching with Comet. **A**, Comet online and offline search scores (XCorr) match exactly, whereas the Comet and SEQUEST search scores deviate similar to previous reports^10^. TMT sample was the TKO interference standard^24^. **B**. Search times for human (blue) and yeast (green) databases. Relative database sizes are depicted as rectangles for each search. Search parameters: forward (F) and reversed (R) decoy peptides, 50 ppm precursor tolerance, default low resolution Comet search, 3 isotopic windows, methionine oxidation [Ox(M)]. **C.** Search times for yeast, human, concatenated human-yeast, and human with STY phosphorylation. For **B** and **C**, the searches were technical replicates of Hyper standard runs^6,23^. Variable modification (VM).

We tested the speed of Orbiter’s RTS using the Hyper two-proteome interference standard^4,6,23^. Searching a full yeast database (6757 protein entries, Uniprot) with a 50ppm precursor tolerance across three isotopes (precursor mass −1/+0/+1) and methionine oxidation as a variable modification and reversed decoy proteins resulted in median search times of 5ms. We next tested Orbiter search times for the significantly more complex full human database with common isoforms (42113 protein entries, Uniprot). When searched with the same parameters as the yeast database, this test resulted in median search times of only 17ms. As noted in the above searches, the RTS revision of Comet (SourceForge revision r1296) retains the flexible database searching available offline allowing the use of user-defined variable modifications, including common post-translational modifications such as methionine oxidation and phosphorylation (Figure 2b). Indeed, we observed median search times for even a full human database considering variable modifications of both methionine oxidation and serine, threonine, tyrosine phosphorylation fell well under the default injection time for 50,000 resolution SPS-MS3 scans (86ms).

In lieu of the E-value calculation (removed to improve search speeds), Orbiter performs a multi-feature linear discriminant analysis (LDA) to distinguish high quality PSMs from low quality PSMs (Figure 3)^9,25^. Orbiter’s LDA uses seven score parameters derived from the output of Comet’s database search: XCorr, deltaCorr, missed cleavages, charge state, absolute ppm error, peptide length, and the fraction of ions matched^15^. The LDA resulted in a final discriminant score that efficiently separated target from decoy PSMs (Figure 3A). Of note, as Orbiter generated a library of PSMs for every run, it was also possible to track and adjust the ppm error on the fly. Thereby Orbiter can actively adapt to the current performance of the instrument (e.g. temperature changes causing Orbitrap mass accuracy to drift). The adjusted precursor mass error measurement also provided a real-time quality control metric to assess aberrant instrument behavior.

**Figure 3.**
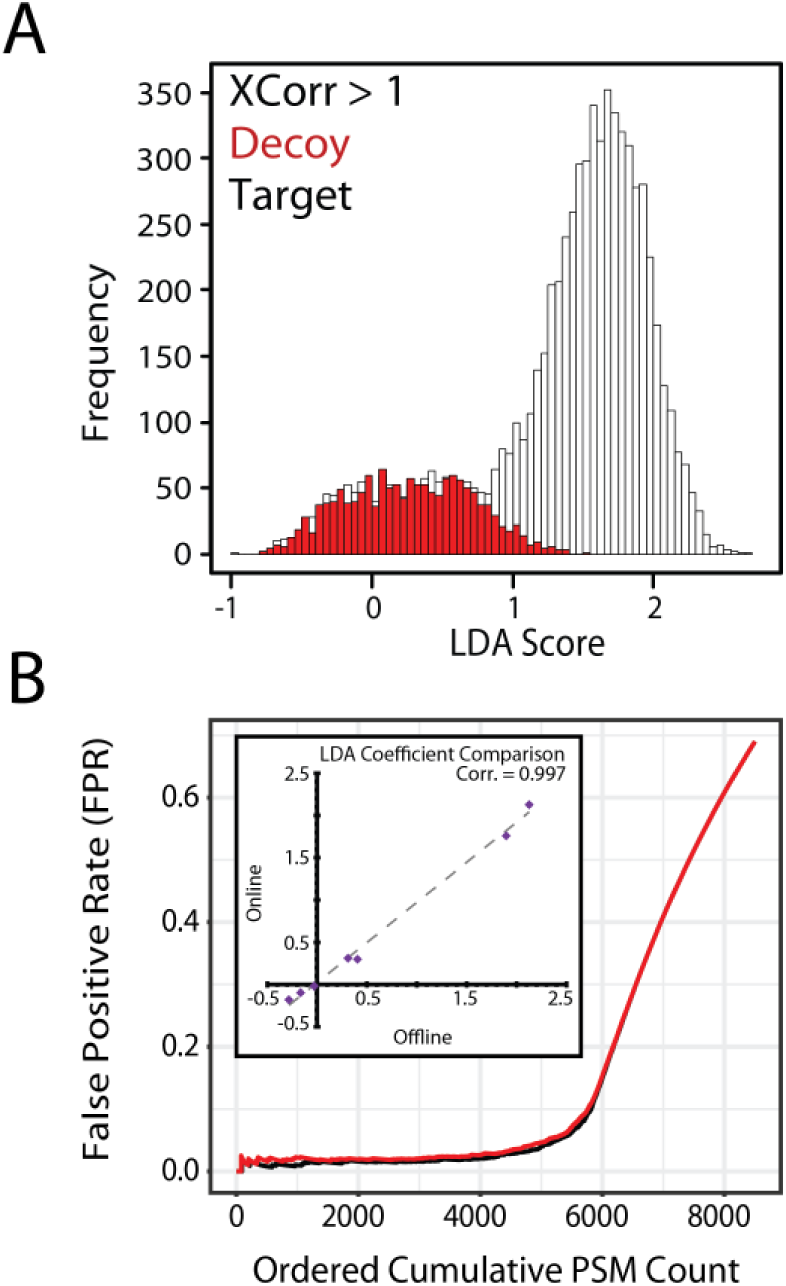
Real-time False Discovery Rate filtering. **A**, Real-time peptide spectral match features (e.g. XCorr, deltaCorr) were used to discriminate target and decoy peptides on-the-fly. **B**, Online and offline LDA scoring resulted in similar numbers of passing peptide spectral matches. Inset: the resulting LDA coefficients from offline and online analysis were highly correlated.

The training data for Orbiter’s LDA was derived from all previous PSMs observed within the single run with and XCorr greater than 1. Therefore, to ensure that this LDA was not biased compared to offline discriminant analysis which used all PSMs from a run for training, we compared online versus offline LDA on the same run (Figure 3B). While the reduced training set did have an effect on the LDA coefficients, this result was minor as noted by the highly correlated LDA coefficients (Figure 3B). Moreover, the resulting discriminant scores across all PSMs were highly similar. When enabled, the PSMs that passed the user-defined FDR filter and did not originate from decoy peptides trigger SPS-MS3 scans. Filtering of low confidence and decoy peptides greatly reduces the number of SPS-MS3 scans triggered. The FDR filter can in turn be combined with protein inclusion and exclusion filters to enable Orbiter to target specific subproteomes, e.g. kinases^4^.

By eliminating SPS-MS3 scans downstream of MS2 scans that matched to low scoring or decoy peptides, Orbiter should improve instrument acquisition efficiency for multiplexed analyses. To assess this, we generated samples from a panel of cell lines of multiple tissues of origin (i.e. mammary gland, colon, embryonic kidney). Biological replicates for each of three cell lines (HEK293T, HCT116, and MCF7) were labeled with TMT reagents and mixed in equal proportion (Figure 4A). The cell line panel was then processed as described previously and pooled into 12 high-pH reversed phase fractions^13^. Each of these fractions was analyzed either with a SPS-MS3 method running 180-minute gradients, or Orbiter methods running shortened 90 minute gradients (Figure 4B). Thus, we tested if Orbiter could reach the same quantitative accuracy and proteome coverage of SPS-MS3 in half the gradient time. After offline filtering to a peptide and protein FDR less than 1%, the SPS-MS3 method quantified 8455 proteins in 36 hours. In just 18 hours, the Orbiter RTS method quantified 8166 (97% of the SPS-MS3 method). These data emphasized a near doubling of the instrument acquisition speed from 234 proteins quantified per hour with SPS-MS3 to 454 proteins quantified per hour with Orbiter RTS. The resulting ratios between SPS-MS3 and Orbiter quantitation were highly correlated (Pearson correlation = 0.946, Figure 4C).

**Figure 4.**
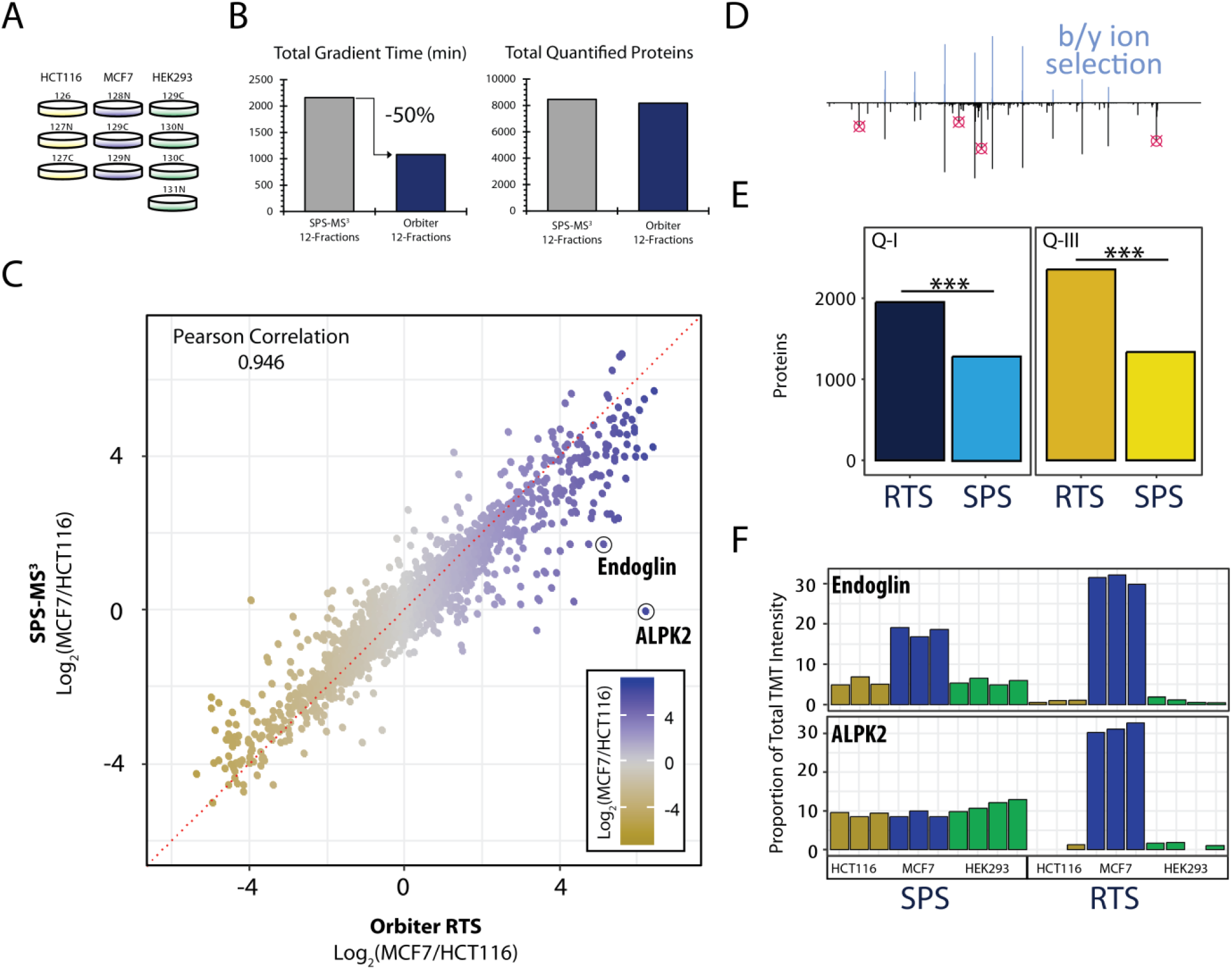
Quantitative comparison of Orbiter’s RTS and canonical SPS-MS3 method. **A**, Cell line panel. **B**, Comparison of total gradient time used and the total number of quantified proteins for each experiment. Orbiter quantified more than 8000 proteins in 50% of the gradient time. **C**, Ratio comparison of Orbiter results and SPS-MS3. The two methods were highly correlated. **D**, the skew seen in C was most likely due to improved quantitation using b- and y-ion selection for SPS ions**. E**, Orbiter’s RTS quantified proteins were significantly enriched for higher quantitative ratios. Quadrants: Q-I (SPS ratio > 0, Orbiter ratio > 0) and Q-III (SPS ratio < 0 and Orbiter ratio < 0). Asterisks (***) denote Fisher’s Exact test p-value < 2.2e-16. **F**, examples of Orbiter’s improved quantitative accuracy for ALPK2 and Endoglin. In both cases, removal of interference markedly improves relative quantitation of these proteins.

Although the ratios were highly correlated, we observed a skew with Orbiter ratios having, in general, greater absolute values (Figure 4C). We built Orbiter’s RTS to select MS3 precursors only from b-ions for arginine terminated peptides or b- and y-ions for lysine terminated peptides (Figure 4D). In contrast, standard SPS-MS3 precursors were selected based on the top-n most intense ions in the preceding MS2 scan. Standard SPS-MS3 ions, therefore, may not originate from the matched peptide^3^. To estimate the extent of the improved RTS accuracy we compared quadrants I and III of the ratio scatter plot to determine if the Orbiter quantification was enriched for larger relative ratios compared to SPS-MS3 with a null hypothesis that an equal number of ratios were greater using Orbiter compared to SPS-MS3 and vice versa. Strikingly, by Fisher’s Exact test we observed a significant enrichment of greater dynamic range measurements using Orbiter (Figure 4E). We highlight two examples of this for the proteins Endoglin (CD105) and ALPK2 (Figure 4F). In each case, Orbiter greatly reduced the quantitative isobaric interference revealing large, cell line specific ratios (Figure 4E).

The Orbiter platform was built as a flexible RTS pipeline that enables rapid deployment of new search-based methods. While the initial use case has targeted improving accuracy and acquisition efficiency in for multiplex-based SPS-MS3 scans, the RTS via Comet could rapidly be extended to diverse applications, such as selection of fragmentation schemes for complex sample types (e.g. ETD or HCD for glycan analysis) based on identifying a specific peptide/post-translational modification or real-time filtering for crosslinking analysis. Accordingly, the core search functionality developed here has been implemented as a new feature for the latest generation of Thermo Tribrid instruments.

